# Multimodal single-cell profiling of T cell specificity and reactivity in lung cancer

**DOI:** 10.1101/2023.10.04.560863

**Authors:** Florian Bieberich, Rodrigo Vazquez-Lombardi, Huixin Jin, Kai-Lin Hong, Petra Herzig, Marcel Trefny, Marta Trüb, Heinz Läubli, Didier Lardinois, Kirsten Mertz, Matthias S. Matter, Alfred Zippelius, Sai T. Reddy

## Abstract

Adoptive transfer of autologous tumor-infiltrating lymphocyte T cells (TILs) offers one of the most promising approaches for cancer immunotherapy. However, high variability in patient responses highlight the need for an enhanced understanding of the transcriptional phenotypes of TILs and reactivity of their T cell receptors (TCR). Here, we employ single-cell multiomics approaches and TCR functional screening to investigate TILs from treatment-naive non-small cell lung cancer patients. This comprehensive analysis integrates scRNA-seq, scTCR-seq, and scATAC-seq, enabling a high-resolution examination of TILs within lung cancer tissue, as well as the adjacent non-tumor tissue. We apply a cellular functional screening platform to identify reactive TCRs that represent >1,000 TILs and have specificity towards a multitude of targets, including primary tumor cells, neoantigens, tumor-associated antigens, and viral antigens. Tumor-reactive TILs were primarily associated with dysfunctional phenotypes, whereas viral antigen-reactive TCRs were found in effector phenotype clusters. Key marker genes were identified and used to construct a tumor or viral reactivity score. Comparing clones shared in tumor and non-tumor tissue, a higher fraction of exhausted cells was observed in the tumor tissue, whereas non-tumor adjacent tissue possessed more effector cells, thus providing insight into potential sources for therapeutic T cells. Elucidating the specific T cell populations within TILs and their associated TCRs may support strategies to enhance the efficacy of TIL-based therapies.

**Graphical Abstract:** *Multimodal single cell profiling and reactivity testing of TILs:* (A) CD8^+^ T cells of treatment naive non-small cell lung cancer patients and adjacent lung tissue were isolated by fluorescence-activated cell sorting (FACS) and were then subjected to scRNA-seq + scTCR-seq or scATAC-seq. (B) TCRs were functionally screened using a cellular platform (TnT cells) and target cells (tumor cells, antigen-pulsed antigen-presenting cells, PBMCs) by flow cytometry and deep sequencing. (C) scRNA-seq + scATAC-seq allowed trajectory inference of transcription factors and genes along pseudotime. (D) Gene scores for tumor- and virus-reactivity were developed by combining functional reactivity and transcriptomic profiling for each CD8^+^ T cell. (E) TIL scRNA-seq pre and post IL-2 treatment in tumor suspension displayed as alluvial plot shows change of clonal cell state composition. 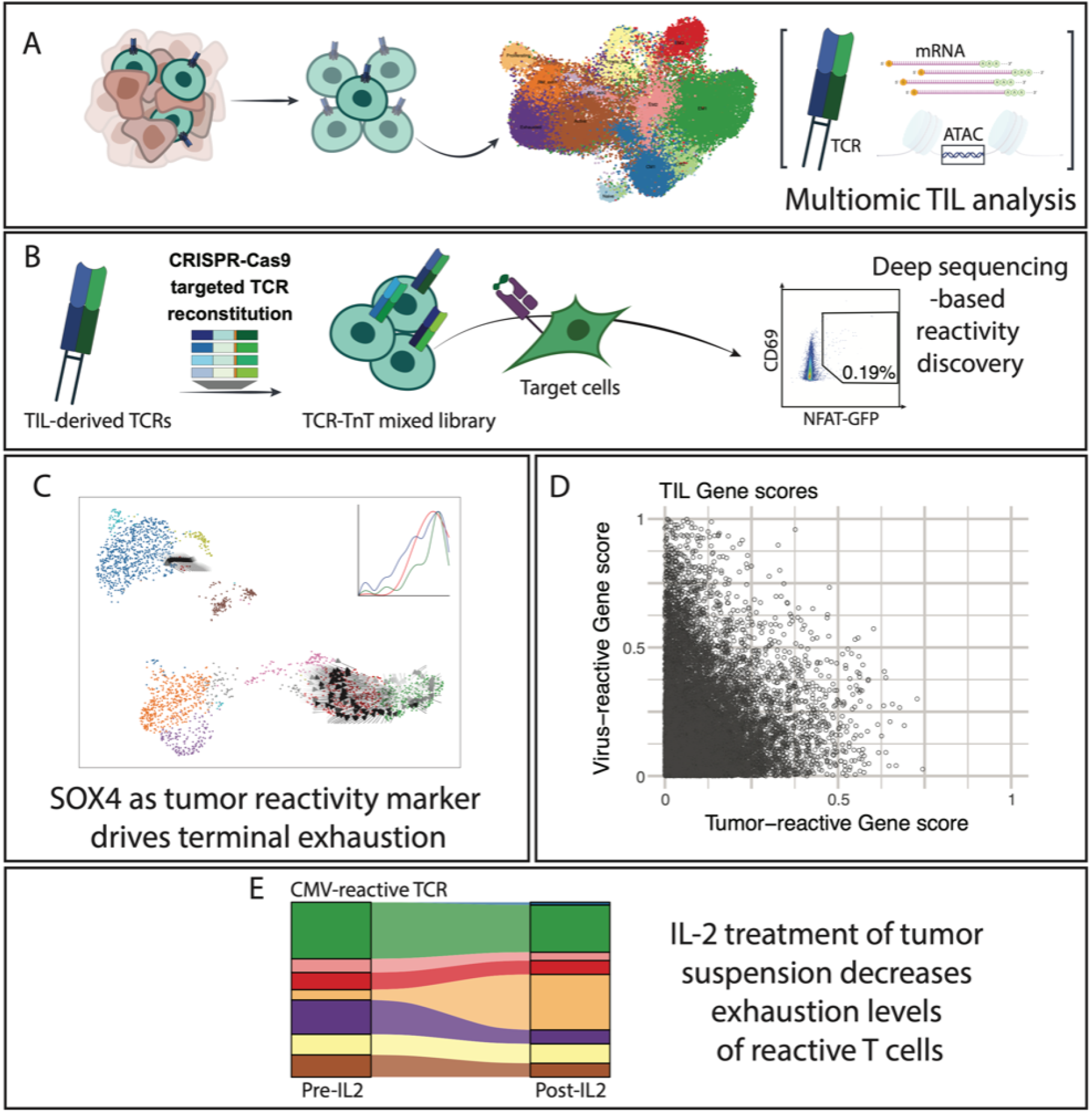

## INTRODUCTION

Lung cancer stands as the primary contributor to cancer-related fatalities worldwide and is characterized ^1–4^ by a high tumor mutational burden, typically exhibiting a substantial amount of immune cell infiltration within the tumor microenvironment ^5–7^. Somatic mutations in the genome of cancer cells leads to the generation of neoantigens, which are mutated proteins that are processed and presented as peptides on human leukocyte antigen / major histocompatibility class I receptors (HLA-I / MHC-I) on tumor cells. Together with tumor-associated antigens (TAA), neoantigens form the core mechanism for tumor cell recognition by cytotoxic CD8 T cells via their T cell receptors (TCR) ^8,9^. Despite these defense mechanisms, a significant proportion of T cells present in a tumor (also referred to as tumor-infiltrating lymphocytes, TILs) exist in a dysfunctional state, rendering them ineffective in halting tumor progression ^10,11,12–15^.

TIL therapy consists of an expanded autologous cell product and when applied for treatment of melanoma has been associated with complete and lasting responses in patients ^16–20^. This outcome is largely due to a subset of tumor-reactive T cells that are capable of driving tumor eradication ^21–23^. However, TIL therapy is only beneficial for a limited number of patients, as often the expansion and reactivity of the TIL products are insufficient, and there is a high prevalence of dysfunctional cell states ^24^. This is likely due to the condition of T cells upon resection and an imbalanced clonal expansion during the expansion phase ^25^, underlining the correlation between T cell phenotypes and the effectiveness of TIL therapy.

T cells reacting to neoantigens provide a highly specific way to target tumor cells while obviating healthy tissue ^26–28^. However, identifying specific peptide-HLA-TCR (pHLA-TCR) pairs is a complex task due to the high diversity of peptides and TCRs within the tumor tissue, as well as the relatively weak (low affinity) interactions between TAA pHLA-TCR pairs ^29–31^. Tumor cells can present a large number of peptides that can potentially be recognized by TCRs, including unique neoantigens and over-expression of TAA, which arise from direct (mutation) or indirect (i.e., post-translational changes) changes in the cancer genome. Besides the large number of potential pHLA-TCR that need to be screened, the low affinity pHLA-TCR interactions pose the need for highly sensitive assays ^32,33^. While computational methods have been used to predict peptide-HLA binding (e.g., netMHC) ^34,35^ and decrease the number of peptides that need to be screened, discovery of reactive TCRs remains challenging ^30,36^. Current methods for TCR reactivity discovery rely on different selection methods, such as binding based detection through T cell labeling with soluble and fluorescently-tagged pHLA multimers ^29,37,38^ or, more recently through co-culturing T cells with antigen presenting cells (APCs) that display tumor antigens through peptide pulsing or minigene expression ^39,40^ and using T cell activation markers or proliferation as readouts ^41–43^. Essential for the high-throughput nature of these assays are TCR repertoire profiling methods such as single-cell and bulk deep sequencing, which facilitate the sequencing of paired (Vα and Vβ) or single TCR chains, respectively ^44,45^. The TCR repertoire is the collection of TCRs present in an individual, tissue, or co-culture setting and can be associated with tumor mutational load and clinical outcome ^46^.

Single-cell RNA sequencing (scRNA-seq) offers the ability to analyze the transcriptional profiles of TILs and combine it with single-cell sequencing of TCRs (scTCR-seq). Innovative methods, such as single-cell multiomics (sc-Multiomics) (simultaneous scRNA-seq and single-cell Assay for Transposase-Accessible Chromatin sequencing [scATAC-seq]), allow for a more detailed assessment of cell state dynamics and regulators ^47^. This involves a combined analysis of transcription factor (TF) transcriptional expression and the genomic accessibility of the TF binding site as T cells progress towards an exhausted state ^48–52^. Consequently, these methods offer a comprehensive insight into TIL cell state dynamics and TCR clonal repertoires and enable functional reactivity assays.

Recent work using single-cell sequencing and functional reactivity testing methods has underlined previous discoveries that neoantigen-reactive TILs are enriched in exhaustion and resident memory cell states. For example, in the studies by Caushi et al. and Lowery et al. it was discovered that only a small proportion of TILs are reactive to neoantigens and selectively express *CD103*, *CD39*, *CXCL13*, *TOX2* and *CXCR6* among others as part of their reactivity signatures ^53,54^. Further, these and similar discoveries by others, have inspired the development of tumor reactivity signatures based on differential expression of various marker genes on both RNA and protein levels ^43,55,56^. TILs often exhibit biased reactivity signaling, which can frequently lead to false negatives in reactivity testing ^57^. Therefore, it is not yet determined what percentage of the actual tumor-reactive TILs can be classified using such molecular signatures, given that most existing gene expression profiles are based on a sparse number of TIL clones. For example, a study by Krishna et al. revealed that patients who responded to TIL therapy retained a pool of stem-like *CD39*-negative TILs, thus contradicting parts of the previously mentioned work and emphasizing the need to explore additional markers of T cell reactivity against tumors ^58,59^.

Here, we profile CD8^+^ TILs from nine patients with primary non-small cell lung cancer resections that had not been previously subjected to systemic therapies (from now on mentioned as treatment naive). We achieve this through a comprehensive single-cell multiomics approach combining scRNA-seq, scTCR-seq and scATAC-seq of TILs from non-small cell lung cancer and non-tumor adjacent tissue. For four patients, we selected a total of 138 TCRs for tumor reactivity testing using a previously established functional cellular screening platform ^60^. T cell reactivity screening was performed against a panel of potential antigens, including common viral antigens, TAAs and neoantigens and led to the identification of 18 reactive TCRs that were represented by transcriptomes of over 1,000 TILs. Further, we identified cell state differences in shared clones in tumor and non-tumor tissue and generated a gene regulatory network that identified drivers of exhaustion. Through short-term IL-2 treatment of tumor suspension we were able to compare the potential cell state change of virus- and tumor-reactive TCR clones, possibly proving valuable for therapeutic decision making. This study combines TIL cell state dynamics with T cell reactivity and provides a helpful resource for the targeted selection of reactive TILs in lung cancer and non-tumor adjacent tissue.

## RESULTS

### CD8^+^ T cell transcriptomes in treatment naive lung cancer patients

Tissue was obtained from nine patients resected for lung tumors who had not undergone therapy prior to resection. Seven patients were males and two females. Patient average age was 67.4 years. Five patients had a squamous cell carcinoma and four an adenocarcinoma (**Table 1, Table S1**).

**Table 1:**
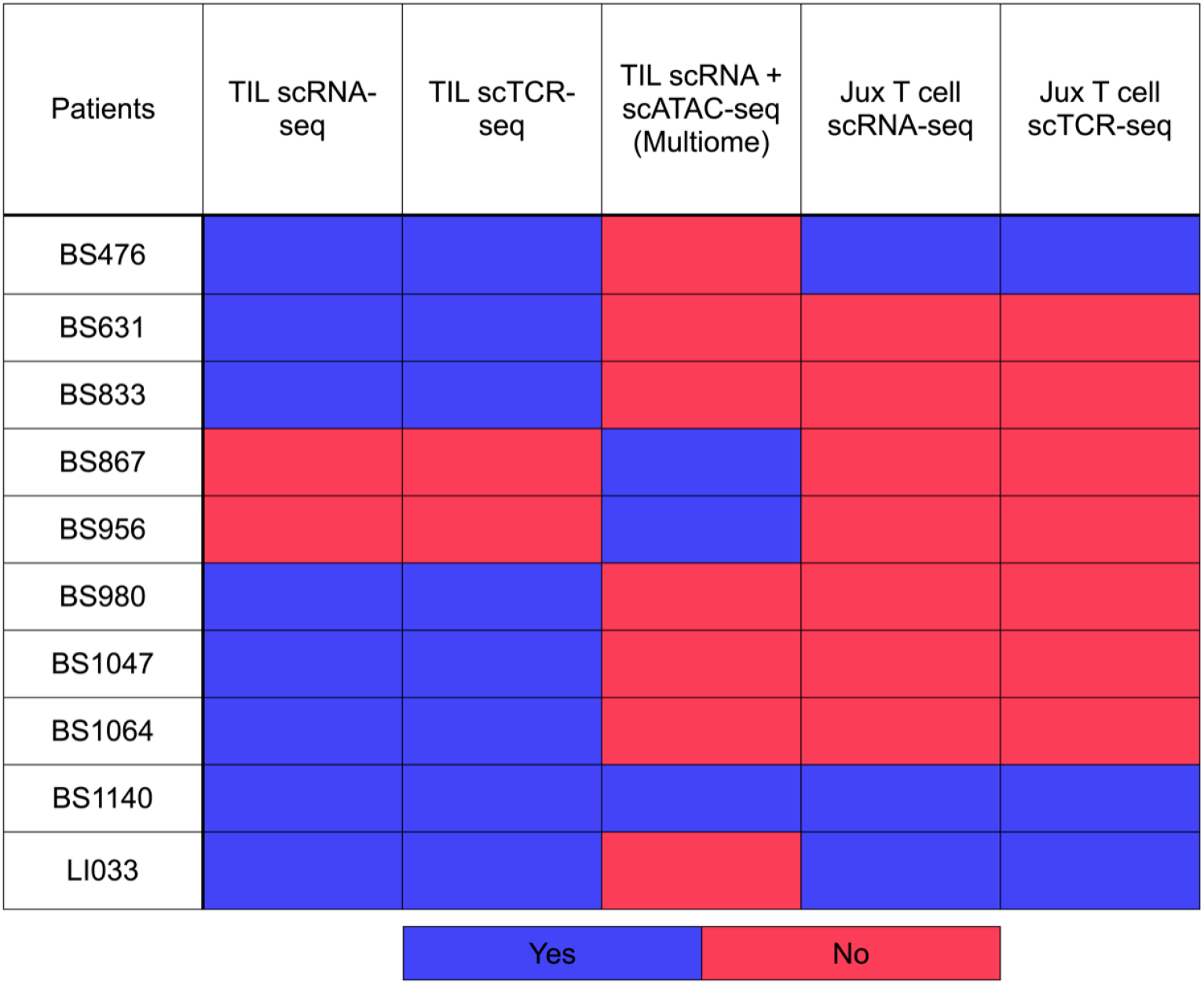
Single cell sequencing methods used in this study for each patient.

We analyzed CD8^+^ T cells (hereafter referred to as TILs) from tumor samples of eight patients, using scRNA-seq and scTCR-seq. For three of the patients (BS476, BS1140 and LI033), T cells from non-tumoral adjacent lung tissue were additionally sequenced for comparative analysis. For another two patients (BS867 and BS1140), TILs were subjected to single-cell multiomics sequencing (scMultiome: scRNA-seq and scATAC-seq from the same cell).

Across all patients and tissue samples, scRNA-seq profiles were generated from 57,432 T cells; and for scTCR-seq, data was generated from 40,228 cells across seven patients. 12 TIL sub-cell states were identified by using a combination of marker genes previously described ^58,61,62^ (**Fig. 1A, B**). Unified manifold approximation and projection (UMAP) analysis was performed and identified exhausted T cell subsets (*Exhausted*, *RM_exh1*) that shared expression of markers such as *CXCL13*, *ENTPD1* (*CD39*) and *PDCD1*, as well as identification of a proximally located “*Proliferating”* cluster associated with *MKI67* expression. Interestingly, expression of innate-like/NK cell markers were present in the *Innate_like_T* and *RM_exh2* cluster, suggesting an NKT-like state next to effector and exhausted cell states. A gene set consisting of markers that were previously shown to correlate with tumor-reactive T cells could be identified in the *Exhausted*, *RM_exh1*, *RM_exh2* and *Proliferating* clusters ^63,64^. A stem-like T cell marker gene set was highly expressed by cells in the *EM2* and *Innate_like_T* clusters, with similar co-location of a recent activation marker gene set. An effector gene set showed broad expression with increased levels in the *Exhausted*, *Innate_like_T* and *EM3* clusters (**Fig. 1D**). Pseudotime analysis indicates that *RM_Exh1* and *Proliferating* clusters contain cells with the most recent gene expression signature and are the most actively cycling cells (similar as shown by Gueguen et al. ^62^) (**Fig. 1E**). Further comparison of cluster frequencies across patients between tumor and adjacent tissue shows that the combined area of clusters *Proliferating*, *RM_exh1*, *RM_exh2* and *Exhausted* is greater than two times as much in TILs from tumor compared to adjacent (non-tumor) tissue, both across patients and by patient pair (**Fig. 1C**).

**Figure 1:**
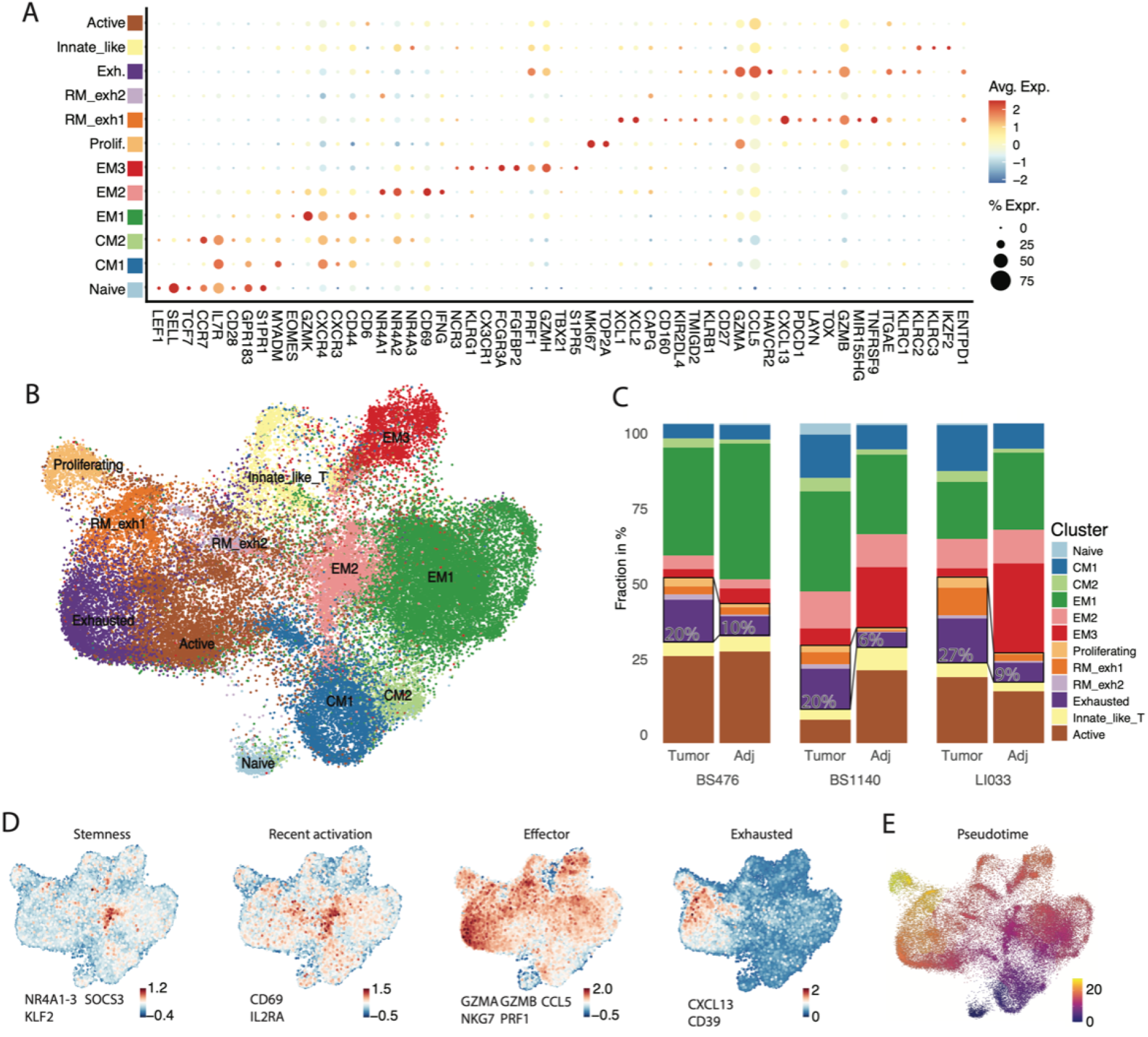
Single-cell transcriptional variability in tumor infiltrating lymphocytes (TILs) from treatment naive non-small cell lung cancer patient samples. (A) Dotplot illustrating marker gene expression across the identified clusters. Color intensity signifies average expression; dot size reflects the percentage of cells within a cluster expressing the marker. (B) Uniform Manifold Approximation and Projection (UMAP) visualization of TIL transcriptome data across multiple patients. (C) Proportion of cells within each cluster derived from tumor and non-tumor adjacent (adj.) tissue. Insets show the cell fraction in clusters: *Proliferating*, *RM_exh1*, *RM_exh2* and *Exhausted*. (D) Module scores calculated using gene sets listed adjacent to each UMAP, indicating average module score expression on a per-cell basis. (E) Pseudotime analysis using the *Naive* cluster as a starting point.

### Identifying reactive TILs by functional TCR screening

Next, we identified TILs that are reactive to tumor-specific mutations as well as to antigens from common viruses, such as cytomegalovirus (CMV) and Epstein-Barr virus (EBV). To achieve this we utilized a recently established TCR functional screening platform based on an engineered human T cell line (TnT cells) ^60^. Briefly, TnT cells were created by immunogenomic engineering of Jurkat cells to introduce *CD8* (*CD8A*, *CD8B*), *Cas9* and a reporter of T cell activation (*NFAT-GFP*), as well as knock-out of beta-2-microglobulin (*B2M*) and the endogenous TCR alpha chain. TnT cells thus serve as cellular recipients for genomic integration of recombinant TCRs, including libraries based on scTCR-seq of TILs. Further, TnT cells provide a functional screening platform for TCR activation by expression of NFAT-GFP or surface expression of CD69 following co-culture with cells (APCs or tumor cells) expressing cognate antigen (pHLA). Compared to primary T cells or TILs, which can have altered cell states and phenotypes that can compromise their activation (e.g., exhaustion), the synthetic TnT cells provide an unbiased functional cell state that can be used for TCR screening, including engineering TCR specificity to target antigens ^60^.

In order to be able to screen a multitude of TCRs and antigens against each other, we leveraged deep sequencing data following selection (FACS) of activated TnT cells that were co-cultured with APCs. To validate our approach, we generated a TCR test library, which included as positive controls two TCR clones (A3WT and A3-05) ^60^ with known reactivity against a TAA of the MAGE-A3 protein (HLA-A*0101-restricted MAGE-A3_168–176_ peptide: EVDPIGHLY) along with an additional 23 TCRs (extracted from patient BS833 and expressed in TnT cells) of unknown reactivity that are not expected to be reactive to MAGE-A3-derived antigens. As target cells, we used the MAGE-A3-expressing EJM tumor cell line or non-MAGE-A3-expressing COLO205 tumor cell line, which both have HLA-A*0101 expression. After co-culture of the TCR-TnT library cells with target tumor cells, FACS was performed to isolate TCR-TnT cells based on the activation markers CD69 and NFAT-GFP (**Fig 2A**). Following sorting, TCR-TnT cells were subjected to targeted deep sequencing of the transgenic TCR locus. Sequencing read TCR frequencies of cells co-cultured with EJM, COLO205 or prior to co-culture (background) revealed a >20-fold enrichment of the control MAGEA3-specific TCRs (A3WT and A3-05) compared to the TCRs with unknown reactivity (**Fig S1**).

**Figure 2:**
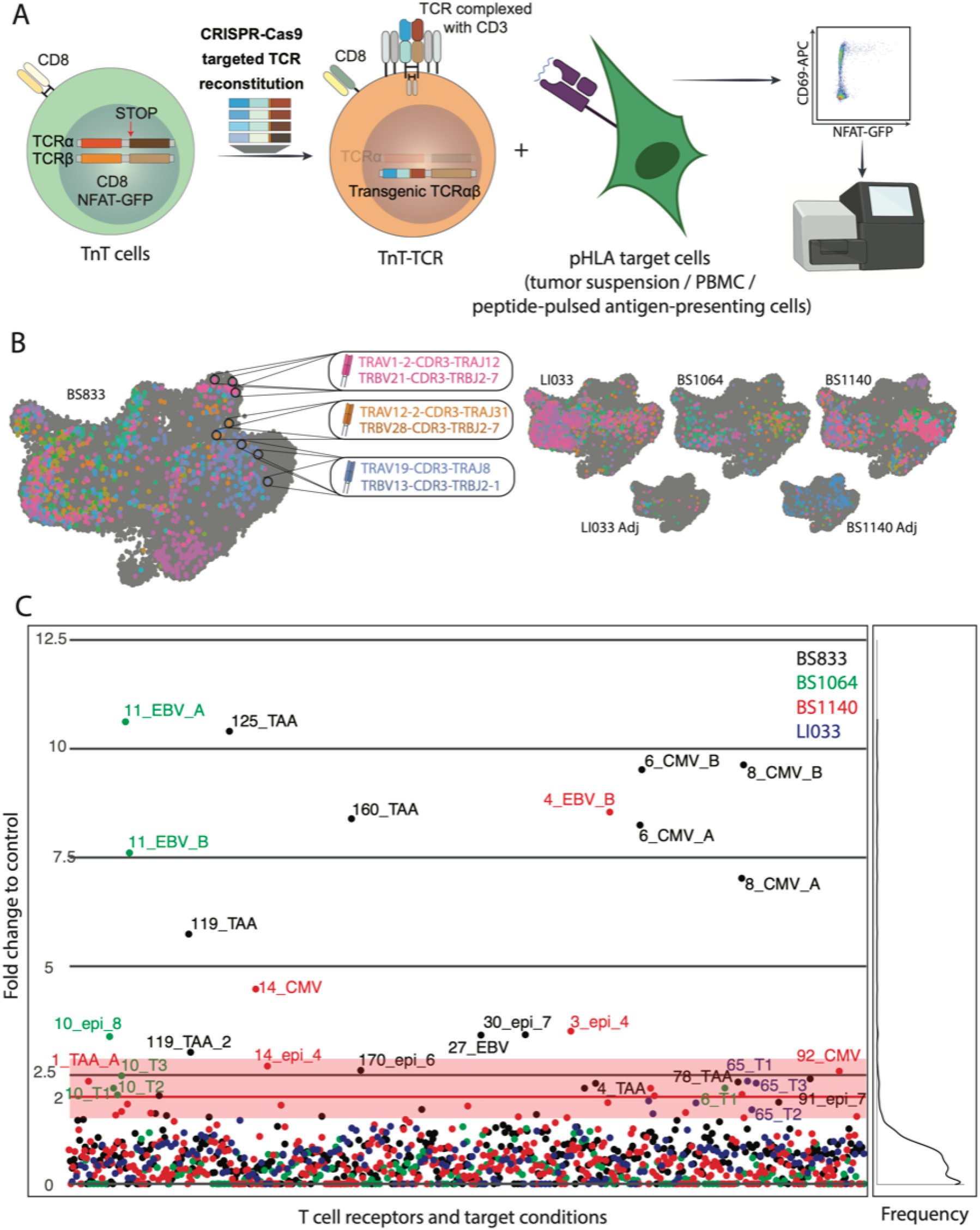
Reactivity profiling of selected TCRs by functional screening and deep sequencing. (A) TCR libraries are genomically integrated into TnT cells by CRISPR-Cas9, which are then co-cultured with pHLA target cells such as tumor suspension, patient PBMC or peptide pulsed antigen presenting cells (see STAR Methods). Fluorescence activated cell sorting (FACS) and deep sequencing are performed to select and identify reactive TCR-TnT cells. (B) T cells with TCR clones selected for functional screening are shown with their respective transcriptomes on UMAP. (C) Left, dotplot shows deep sequencing results following TCR-TnT functional screening. Fold-change represents TCR sequencing reads of each target reactivity condition compared to the highest control value, across all control conditions for the same TCR, color corresponding to the patient. Reactive TCRs are designated as those with fold-change values >2.5. Right, density plot of deep sequencing results for all TCRs and their target conditions across patients. X-axis represents the frequency of all TCR-condition values per fold-change value that is shown as in (left) on the y-axis.

Leveraging scRNA-seq and scTCR-seq data, we rationally selected TCR clones to screen in TnT cells. Here, we define TCR clones as T cells that have identical V(D)J germline usage and have 100% amino acid sequence identity of CDR3α and CDR3β sequences. The TCR clones selected for screening were based on several parameters that may be associated with reactivity: i) highest clonal abundance in a given patient, ii) the ten most abundant clones in each of the following clusters: *Proliferating*, *Exhausted, RM_exh1 or RM_exh2*, iii) clones that were co-abundant in both *EM1* and one or more of the previously mentioned clusters (see **Methods**; **Fig 2B**). The selected TCRs were integrated in TnT cells by CRISPR-Cas9 homology-directed repair (HDR) and the resulting TnT-TCR libraries were screened as a pool with co-cultures of autologous tumor cell suspension or co-culture with patient-specific HLA-presenting cells that were pulsed with either neoantigens, multiple TAA linked to T cell reactivity in lung cancer (polyTAA: *WT1/WT33*, *CEA*, *NY-ESO-1*, *MAGE-A3*) ^65^ or viral antigens derived from EBV and CMV. Neo-antigens were selected based on in-silico predicted binding strength of mutated peptides from exome sequencing data to patient-specific HLAs (see **STAR Methods**). TnT-TCR cells co-cultured with patient-derived PBMCs that did not receive any peptide were used as a negative control. The bacterial superantigen phytohemagglutinin (PHA) was used as a positive control to detect nonspecific activation.

TCR reactivity data for identification of cognate pHLA was generated as follows. FACS was used to isolate TCR-TnT cells expressing above-background levels of the dual activation markers *NFAT-GFP* and *CD69* (**Fig S2**). Next, targeted deep sequencing was performed on the transgenic TCR regions of TnT cells. Deep sequencing data was analyzed to determine TCR clones that were enriched under the different co-culture conditions. TCR clone frequencies were compared between target and control co-culture conditions (pre co-culture, PBMC co-culture, PHA-activated co-culture). Through statistical analysis, each TCR clone per target co-culture received a fold change in comparison to the highest control co-culture. A reactivity threshold could be determined based on the density plot of fold-change values across all TCR clones and target co-cultures. TCR clones with a fold-change value > 2.5 were classified as reactive (see **Methods**; **Fig S3; Table S2**).

This led to the identification of 18 reactive TCRs (out of 138 tested TCRs). Multiple TCRs showed similar fold change reactivities for each duplicate against CMV (BS833, clone 6 and 8), EBV (BS1064, clone 11) and pTAA (BS833, clone 119), showing the robustness of this functional screening approach. One TCR showed cross reactivity between neoantigen/self-antigen and CMV (BS1140, clone 14); with another one showing reactivity against tumor suspension and neoantigen (BS1064, clone 10) (**Fig 2C**). Genes that harbored mutations for neoantigen generation and were associated with TCR reactivity were mutated in 2.5%-10% ^66–68^ of lung cancer patients and included: *KMT2D*, *ATR* and *SORCS1*.

### Phenotypes of reactive T cells in treatment naive lung cancer biopsies

Following identification of 18 TCRs with reactivity to tumor cells and viral antigens, we next analyzed the associated transcriptional phenotypes that consisted of over 1,000 corresponding TILs. Tumor- and polyTAA-reactive TILs showed high abundance in clusters associated with (pre-) dysfunctional phenotypes (**Fig S5**), whereas CMV and EBV viral antigen-reactive TCRs were mainly present in effector phenotype clusters (**Fig. 3A**). Patient tissues showed absence of expression of EBV or CMV, indicating that these virus-reactive TCRs respond to systemic antigens rather than tissue-specific antigens (not tumor-specific) (**Table S3**). We subsequently determined key marker genes in TILs associated with tumor or viral reactivity by differential gene expression analysis between these two subgroups. The strongest marker for tumor reactivity is *CXCL13*, which corroborates a previous study ^64^ (**Fig. 3B**). *SOX4* is a transcription factor that facilitates the development of *CXCL13*-producing T cells ^69^ and is also among the five genes with the highest differential gene expression for tumor reactivity, further suggesting the importance of the *CXCL13* pathway for TIL activation. Other tumor reactivity markers included genes involved in co-stimulation (*CD7)* ^70^, repeated antigen stimulation (*DUSP4)* ^71^ and immune checkpoints (*TIGIT)* ^72^. For virus reactivity, cytotoxic markers such as *GZMK* and *GZMH* showed high abundance in EBV- and CMV-reactive TILs, respectively. Analysis of T cell reactivities across clusters revealed a strong enrichment of tumor reactivity in the *Exhausted*, *RM_exh1*, *RM_exh2* and *Proliferating* clusters. It is important to consider that since non-reactive T cells showed abundance in clusters that were enriched for CMV-reactive and tumor-reactive T cells, they may still possess reactivity to antigens that were not included in the co-culture screening (**Fig 3C**).

**Figure 3:**
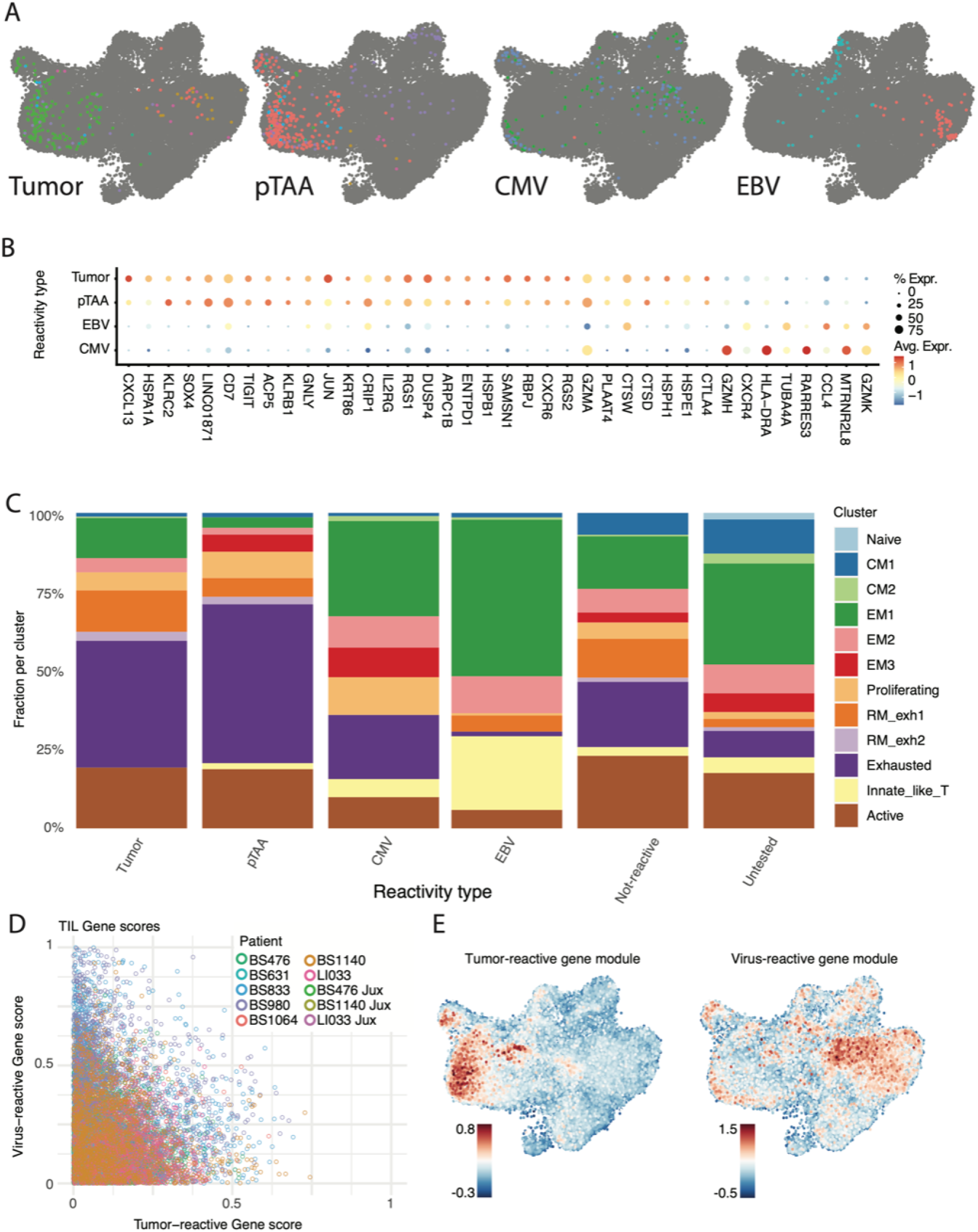
Characterisation of transcriptional programs of reactive TILs. (A) Reactive T cells highlighted on the UMAP according to target tumor or virus antigens. Tumor includes reactivity against tumor suspension and neoantigens; polyTAA (pTAA) includes multiple TAA: *WT1/WT33*, *CEA*, *NY-ESO-1*, *MAGE-A3*. (B) Dotplot map showing each identified marker gene for tumor and virus reactive T cells. Rows correspond to the reactivity groups of T cells. Color indicates average expression, dot size indicates fraction (%) of cells in a reactivity group. (C) Fraction of cells per cluster sorted by each reactivity group of T cells. Non-reactive T cells are defined as TCRs that were functionally screened but did not show reactivity. Untested are all TCRs that were not subjected to functional screening. (D) Scatter plot of tumor- and virus-reactive gene scores, whereby each dot represents a cell and is color-coded according to patient and tissue. Gene scores were derived by combining differentially expressed gene expression levels (see STAR Methods). (E) Tumor (left) and virus (right) gene module score expression on UMAP; score range is relative to each gene module score.

To infer likelihood of reactivity in untested cells, we developed a reactivity score based on gene expression similarity to the experimentally validated tumor- and virus-reactive TILs. This is accomplished by creating a gene module score, which is a linear combination of gene expression levels, derived from the differentially expressed genes observed in tumor- or virus-reactive T cells. (**Fig 3D, Fig S6**). When visualized on the UMAP, these two module scores show *Proliferating*, *Exhausted* and *RM_exh1-2* or *EM1-3* as enriched areas for locations of tumor- or virus-reactive T cells, respectively (**Fig 3E**).

In agreement with previous studies that showed little to no correlation between T cell clonal expansion level and detected reactivity ^32^, we did not observe enrichment of reactive TCRs in the most expanded clones (**Fig S7**).

### TCR clonal sharing in treatment naive lung cancer and non-tumor adjacent tissue

The combination of scRNA-seq and scTCR-seq is able to connect transcriptomes with full TCR sequences (paired Vα and Vβ) and thus elucidate cell state diversity across T cell clones. From the 40,228 cells with scTCR-seq data, 26,018 had corresponding scRNA-seq data (**Fig S8**). Among these, 12,284 scRNA-seq and scTCR-seq cells were in patients with both tumor (8,854) and adjacent (3,430) tissue (**Fig 4A**).

**Figure 4:**
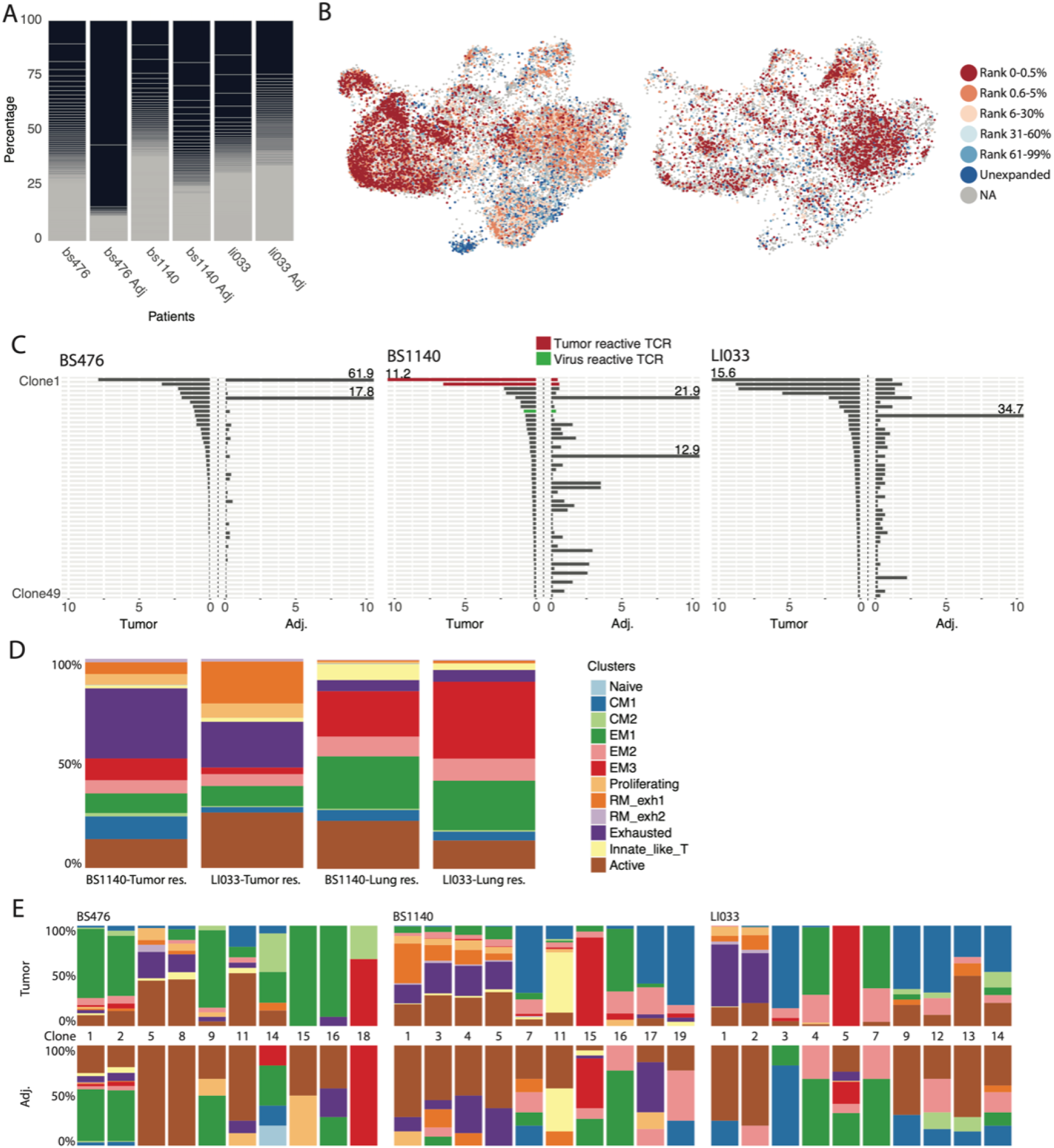
Differential TIL phenotype of shared clones in tumor and non-tumor adjacent lung tissue. (A) The TCR clonal distribution as percentage of the observed repertoire. Grey lines separate clones. (B) Clonal expansion levels for TCRs mapped onto the UMAP. Left, TCRs of T cells that were identified in patient tumor tissue. Right, TCRs of T cells that were identified in patient non-tumor adjacent (adj.) lung tissue. Color represents clonal fractions according to their expansion level. Unexpanded clones are defined as TCRs found in only one cell. (C) The 49 largest clones shared between tumor (left) and adjacent (adj.) (right) tissue. Each line represents one clone. X-axis shows the percentage of TCRs in the corresponding patient repertoire. If the percentage exceeded the x-axis limits, the number indicates the clonal percentage. Lines are colored by reactivity to tumor (red) or virus (green). (D) Cluster distribution of all T cells that are resident by majority in tumor or adjacent lung tissue by patient, according to the previous tissue abundance measurements. (E) Transcriptome of the 10 largest shared clones across both tissue modalities. Shown as percentage of clone per cluster.

The scTCR-seq repertoire analysis revealed major differences in T cell clonal expansion levels across tissues. T cells from adjacent tissue show larger clonal expansion levels and in turn also reduced diversity of clones compared to T cells in the tumor counterpart, especially for the most expanded clones (**Fig 4A**). Across TIL TCR repertoires, the largest five clones accounted for more than 25% of the cells. Expansion level, based on clonal size, describes the number of cells identified per clone and can be caused by extensive proliferation through TCR activation upon cognate pHLA binding. Transcriptome analysis associated with clonal expansion levels (here, shown as percentage of expanded clones that is ordered by rank of clonal size) revealed preferential phenotypes between T cells in tumor and adjacent (non-tumor) lung tissue. In the tumor tissue, unexpanded clones (1 cell per clone detected) are mostly located in the *Naive* and *CM1*-*2* clusters, and moderately expanded clones (*Frequency 6-99%*) are evenly distributed across *CM1-2*, *EM1-3*, *Exhausted* and *Proliferating* clusters. Whereas the highest expanded clones (*Frequency 0-5%*) show enrichment in the *Exhausted*, *RM_exh1*, *RM_exh2*, *Proliferating* clusters (**Fig 4B, left**). Interestingly, the highest expanded clones that reside in the adjacent tissue are almost exclusively located in the *EM1-3* clusters (**Fig 4B, right**). Compared to the tumor tissue, this shows vastly different T cell phenotypes for cells with the same expansion levels. To discern which TCRs are predominantly located in the tumor as opposed to the adjacent non-tumor lung tissue, we contrasted the frequencies and cellular states of matched TCRs (clones observed within both the tumor and adjacent tissue) (**Fig S9**). The largest matched clones identified were generally found in greater abundance within the tumor. However, among the top 100 matched clones, over 10% appear to be in greater abundance within the surrounding tissue, suggesting potential circulation from the adjacent tissue into the tumor. Conversely, clones larger in the tumor may be migrating outward into the adjacent tissue (**Fig 4C**). This T cell residency is also reflected in a differing phenotype that could be identified when comparing cell states of T cell clones that were more than 3-fold enriched in one over the other tissue type (tumor- or lung (adjacent)-residency). There was a marked increase of effector T cell states when contrasted with the resident memory, exhaustion, and proliferating cell states found in tumor-resident T cells (**Fig 4D**). Further analysis of the 20 largest shared clones across tissues revealed higher instances of exhaustion in the clones present in the tumor. This suggests the adjacent lung tissue could be a more likely source of effector-like T cells (**Fig 4E**).

### Single-cell multiomic analysis reveals new drivers of a T cell terminal exhaustion phenotype

The state of T cells is significantly influenced by their environment and the stimuli they receive through cytokines and TCR signaling. To study the dynamics of T cell states in untreated, primary human lung adenocarcinoma, we used a sc-Multiomic analysis that integrates scATAC-seq with scRNA-seq. This approach allowed us to study both chromatin accessibility and gene expression in the same cell (**Fig 5A**). With scATAC-seq, it is possible to identify DNA motifs that act as enhancers or repressors through TF binding. scMultiomics also provides gene expression measurements of TF-encoding genes, TF-target genes and accessibility of TF-motifs; the combination of these three features is referred to as a Regulon ^73^. Regulons were identified across all T cells and used for a new clustering analysis, resulting in 10 clusters (**Fig 5C**). Regulon analysis also revealed a major difference in clustering of exhausted and proliferative cells (*clusters 2* and *3 in Fig 5C*) in comparison to using only the scRNA-seq or scATAC-seq data (**Fig 5B, Fig S10, Fig S11**). Consistent with the previously shown analysis (**Fig 1E**), pseudotime in the Regulon UMAP displays exhausted and proliferating clusters as the latest cells along the trajectory (**Fig 5F**). Further, analysis of TCRs from sc-Multiomic sequencing shows comparable transcriptomes for the same TCR clone in scRNA-seq data (**Fig S12**). Differential Regulon analysis identified multiple known TFs implicated in specific T cell states such as *LEF1* and *BACH2* in the naive, *EOMES* in the effector and *PRDM1* in the exhausted state. We also identified multiple TFs that are uniquely implicated in the exhausted state such as *SOX4*, *RFX2*, *ETS1*, *ETV1* and *STAT1* (**Fig 5D**). To contextualize Regulons within cellular differentiation, we constructed a gene regulatory network (GRN) ^74^. We marked TFs and their encoding genes with names that are associated with certain cell states or that have a high connectedness in the network (centrality). The lower area shows enrichment of genes implicated in exhausted T cells, such as *TOX*, *CD39* and *CXCL13*. Genes that cluster together with such may have an effect on T cell persistence and exhaustion (**Fig 5E**). For example, it was just recently shown that *SNX9* deletion alleviates T cell exhaustion ^75^. Detailed analysis of TFs that are strongly associated with driving exhaustion (*RUNX1* and *PRMD1*), as well as being a marker for tumor-reactive TILs (*SOX4*) visualizes their impact on T cell differentiation towards the exhausted state (**Fig 5G**). In line with this, markers represented in the tumor-reactive signature that are represented in the GRN are clustered almost uniquely in the exhausted area (**Fig 5E**). While TF expression and target gene expression steadily rise along the exhaustion trajectory, the target region accessibility abruptly increases in the last exhaustion stage, possibly corresponding with the transition of early (reversible) to late (irreversible) dysfunction (**Fig 5G**).

**Figure 5:**
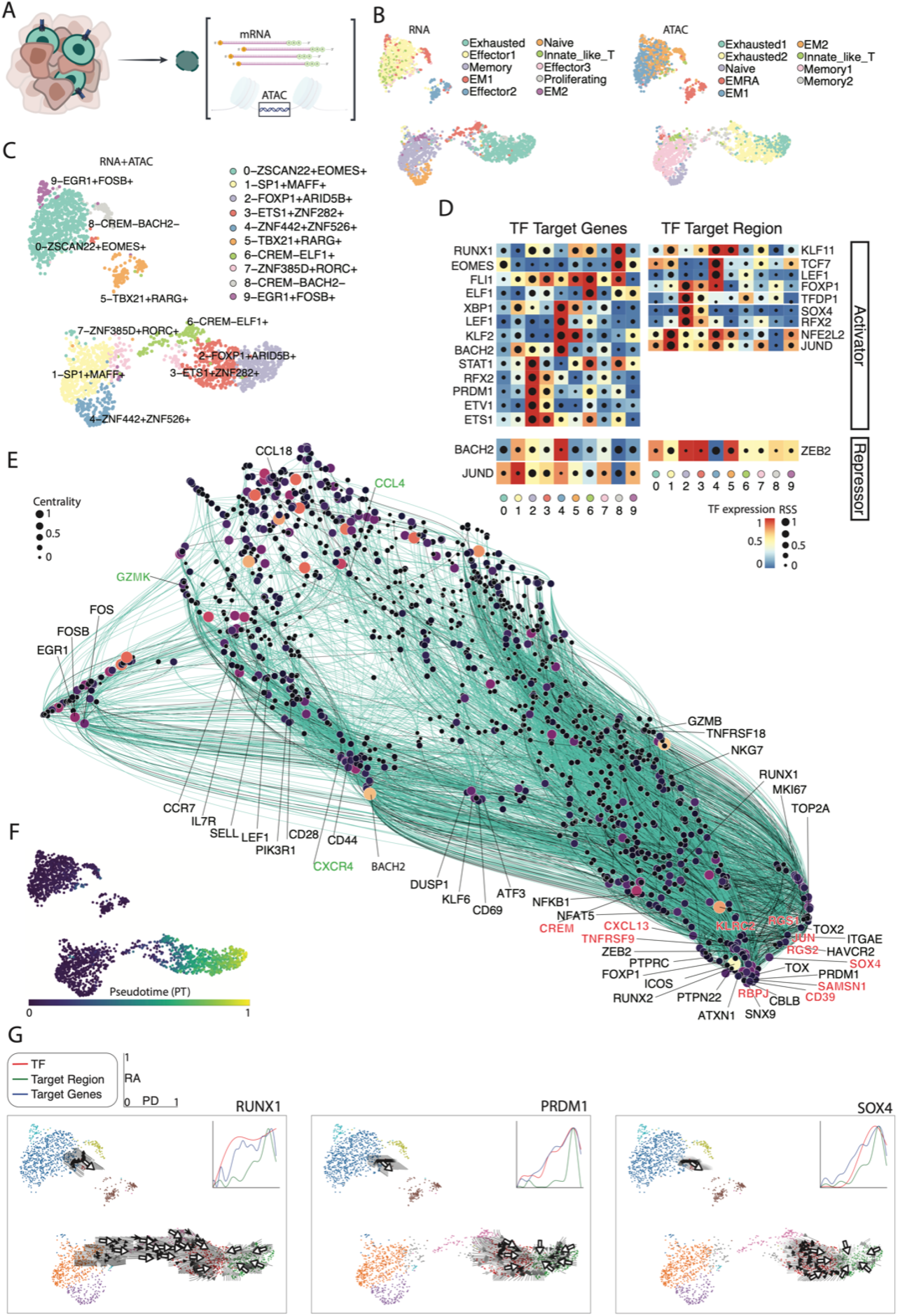
Applying scMultiome sequencing for gene regulatory network analysis of TILs. (A) Schematic of workflow for scMultiome sequencing (scRNA-seq and scATAC-seq). (B and C) Clustering was done via SCENIC+. (B) Left, UMAP cluster coloring based on scRNA-seq data alone. Right, UMAP cluster coloring based on scATAC-seq data alone. (C) Multiome UMAP cluster coloring based on Regulon information for each cell. Regulon is a combination of scATAC- and scRNA-seq data for TF expression, TF target-gene expression and TF-binding site accessibility. Each cluster was named by the most differentially expressed and active Regulon. (D) Differentially expressed transcription factors (TFs) and their target genes (RNA-based) (left) or target regions (ATAC-based) (right) according to the scMultiome UMAP. (E) Gene regulatory network (GRN) using Regulon-information across all cells from patient BS1140. Color and size correspond to centrality (large and bright = high centrality). Line color indicates enhancing (turquoise) or repressing (grey) interaction between genes. (F) Pseudotime of Multiome UMAP. (G) Arrow- and Curve-plots of TFs. Arrow plots indicate differentiation force and direction, based on scMultiome-seq data. Curve plots show TF expression level (red), target gene expression level (blue) as well as specific region accessibility (green) along the pseudotime (Fig 5F) from naive (cluster 4) to exhaustion (cluster 2 and 3).

### Tumor suspension IL-2 co-culture leads to a decreased exhaustion cell state in tumor- and virus-reactive TILs

Having discovered gene reactivity markers and clones with associated reactivities that are also present in the *Proliferating* cluster, we determined how these clonal phenotypes change in short-term (2 days) culture conditions in order to simulate tumor suspension-based stimulation of reactive T cells. We thus chose normal concentrations of IL-2 (100 IU/mL for 2 days) as the culture condition of TILs and used a tumor single cell suspension from patient BS833 that contained TIL clones with various reactivities. Analysis of scRNA-seq from TILs, both pre- and post-IL-2 treatment, revealed a 4-fold larger fraction of TILs in the *Proliferating* cluster; crucially, there was a stronger exchange of RNA velocity vectors between the *RM_exh1* and *Proliferating* clusters, suggesting active transfer between these two cell states (**Fig 6A**). Comparison of reactive T cells from pre- and post-IL2 treatment showed clear enrichment of virus-reactive (CMV) and neoantigen-reactive T cells in the *Proliferating* cluster. Neoantigen-reactive T cells are driven into an *Innate_like_T* cell state and reduce their *Exhausted* cell state upon IL-2 treatment (**Fig 6B**). The 20 largest shared clones in pre- and post-IL-2 treatment data showed overall increases in *Proliferating*, *Innate_like_T* and *EM1* states and decreases in *Exhausted*, *CM1-2* and *EM2* states post-IL-2 treatment (**Fig 6C**).

**Figure 6:**
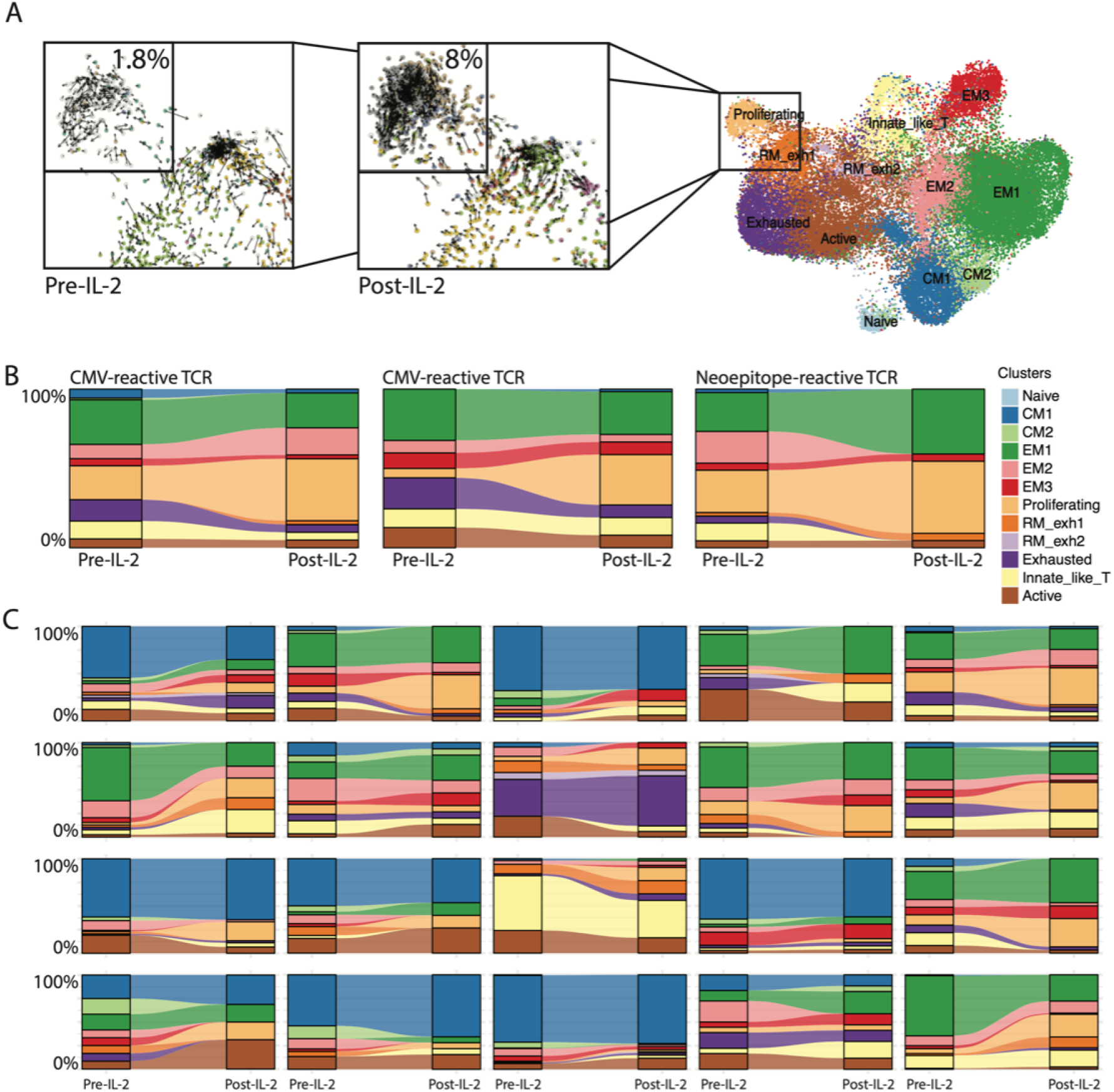
TIL cell state changes upon low IL-2 treatment while in tumor suspension. (A) UMAP showing transcriptome data of cells of patient BS833. Zoom windows show RNA velocity representations of the *Proliferating* cluster pre- and post-IL-2 treatment. The smaller window represents the percentage of cells being part of the *Proliferating* cluster for both timepoints. (B and C) Alluvial-plots showing the change of cell-state distribution per T cell clone pre- and post-IL-2 treatment as percentage of clusters. (B) Shown are three TCR clones with known reactivities. (C) Shown are the 20 largest shared clones for pre- and post-IL-2 treatment data (from left to right and top to bottom).

## DISCUSSION

In this study, we profiled CD8^+^ TILs from untreated primary lung cancer patients by using single-cell multiomics, which combined scRNA-seq, scTCR-seq, and scATAC-seq and was followed by functional screening to identify reactive TCRs.

Importantly, by using a previously established TCR functional screening platform based on an engineered human T cell line ^60^, we were able to avoid the heterogeneity of cellular phenotypes present in primary TILs, which are often used for screening endogenous or transgenic TCRs ^43,54^ and can be a source for confounding results. For example, dysfunctional TILs upregulate CISH, a key regulator of T cell functional avidity, thus leading to an increased activation threshold upon cognate pMHC binding and in turn this may result in higher false negative rates of TILs in co-culture screens ^57^. While functional screening was able to identify reactive TCRs, it may still be possible that some tumor-reactive TILs are already dysfunctional at the time of scRNA-seq and thus did not upregulate gene activation markers of proliferation upon cognate (tumor-) antigen binding. Furthermore, the selection of antigens for reactivity testing is critical for interrogation of the tumor-reactive TIL population. Therefore, we included a diverse mix of neoantigens, TAA, and patient-derived tumor cells. While other studies have primarily focused on neoantigens (minigenes expressing proteins with tumor-specific mutations identified by whole exome sequencing) ^53,54^, their sole use may underestimate the number of reactive TILs ^76^. In future studies, it may be beneficial to couple TCR reactivity screening with de-orphaning of antigen specificity through high-throughput pHLA antigen screening systems ^77–80^.

Our study identified a distinct transcriptomic signature in tumor-reactive T cells. This signature highlights gene expression patterns linked to repeated antigen stimulation ^61,81^, emphasizing its connection to recently discovered markers of exhaustion such as SNX9, as revealed by our GRN ^53,75^. Central to this tumor reactivity signature is the exertion of immune cell recruitment via the CXCL13 pathway, including SOX4, and the upregulation of immune checkpoints (e.g., ITGAE), T cell differentiation markers (e.g., CD7 and DUSP4), as well as a distribution among clusters of T cell exhaustion, stemness, resident memory and proliferation ^69–72^, possibly being the result of asymmetric cell division upon antigen encounter ^82,83^. Notably, CXCL13 emerged as a robust marker for tumor-reactivity, corroborated by findings in other tumor types and has also been linked to tertiary lymphoid structure formation ^54,64,84,85^. Furthermore, markers such as CD7 and DUSP4, suggestive of repeated antigen stimulation, alongside resident memory marker ITGAE, underscore the heightened interactions between T cells and tumor cells within the tumor microenvironment .

The importance of T cell state of stemness for TIL adoptive cell therapies has been demonstrated previously ^58^, however neoantigen-reactive TILs have been observed to lack such stemness phenotypes ^43,53^. Here, we find a substantial overlap of the tumor-reactive gene module and stemness in TILs, possibly owing to our inclusion of polyTAA and tumor suspension as antigen targets, as well as focusing on treatment naive non-small cell lung cancer patients. Interestingly, for shared TCR clones (cells present in tumor and the non-tumor adjacent tissue), the T cells from adjacent tissue display lower levels of dysfunction, indicating a bifurcation of T cell states of the same clonal origin, possibly through lowered antigen exposure or suppressive signal in the adjacent tissue. Here, we observed a significant clonal T cell overlap between tumor and non-tumor adjacent tissues, corroborating previous findings ^62^. Moreover, through our functional reactivity analysis, we determined that tumor-reactive T cells not only migrate from the tumor to the adjacent non-tumor tissue but also exhibit a pronounced effector-like phenotype within this adjacent milieu.

In addition to the identification of a tumor-reactive signature, the scMultiomic approach may also support the preferential enrichment of tumor-reactive T cells through a gene network of exhaustion markers, likely caused by repeated antigen stimulation. The identification of TCRs could be used or engineered for molecular or cell therapies ^86–88^. Recent findings demonstrate that single TCRs could be harnessed as therapeutics for multiple cancer types by exploiting their cross-reactivity against multiple tumor antigens ^89^; this underlines the importance of identifying tumor-reactive TCRs, potentially through methods such as gene signature enrichment.

Moreover, using non-tumor adjacent lung tissue as an additional source for reactive T cells may enable the extraction of more effector-like tumor-reactive T cells, which may help with expanding a sufficient number of T cells for autologous therapy ^90^. As IL-2 is an integral part of many TIL-rapid expansion protocols, we could also show that short-term culture with low-dose IL-2 and tumor suspension leads to varied changes of clonal cell state diversity and could serve as a predictor for patients’ response to immunotherapies, as well as offering potential new strategies for IL-2-based expansion ^91^ or targeting of T cells ^92^. Long-term, it will be critical to understand the relationship between TIL cell states and therapeutic outcome, as is currently being done for chimeric-antigen receptor T cell therapies ^93^.

## SUPPLEMENTARY MATERIALS

Figs. S1 to S11

Tables S1 and S2

## Methods

### Key resources table

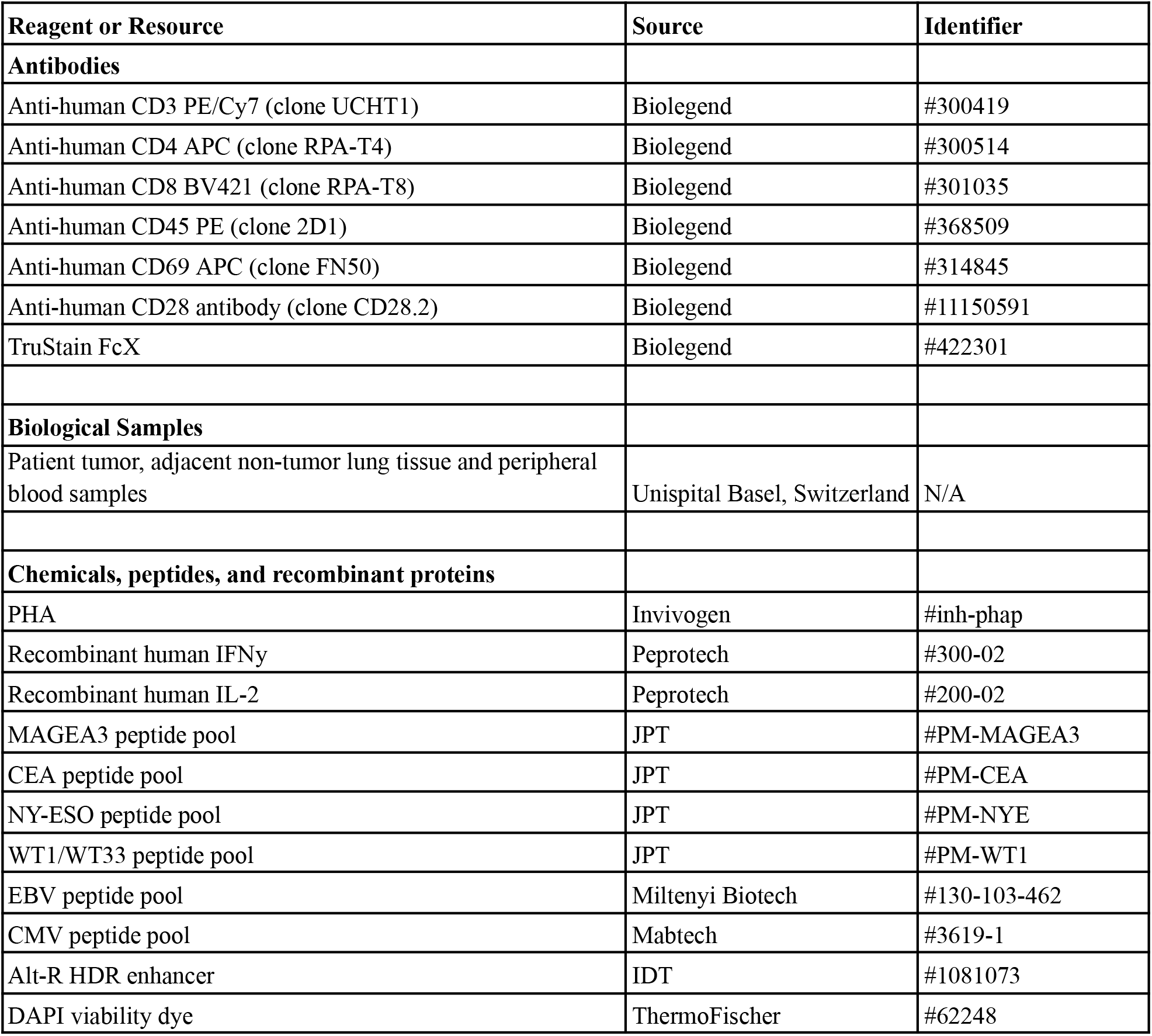
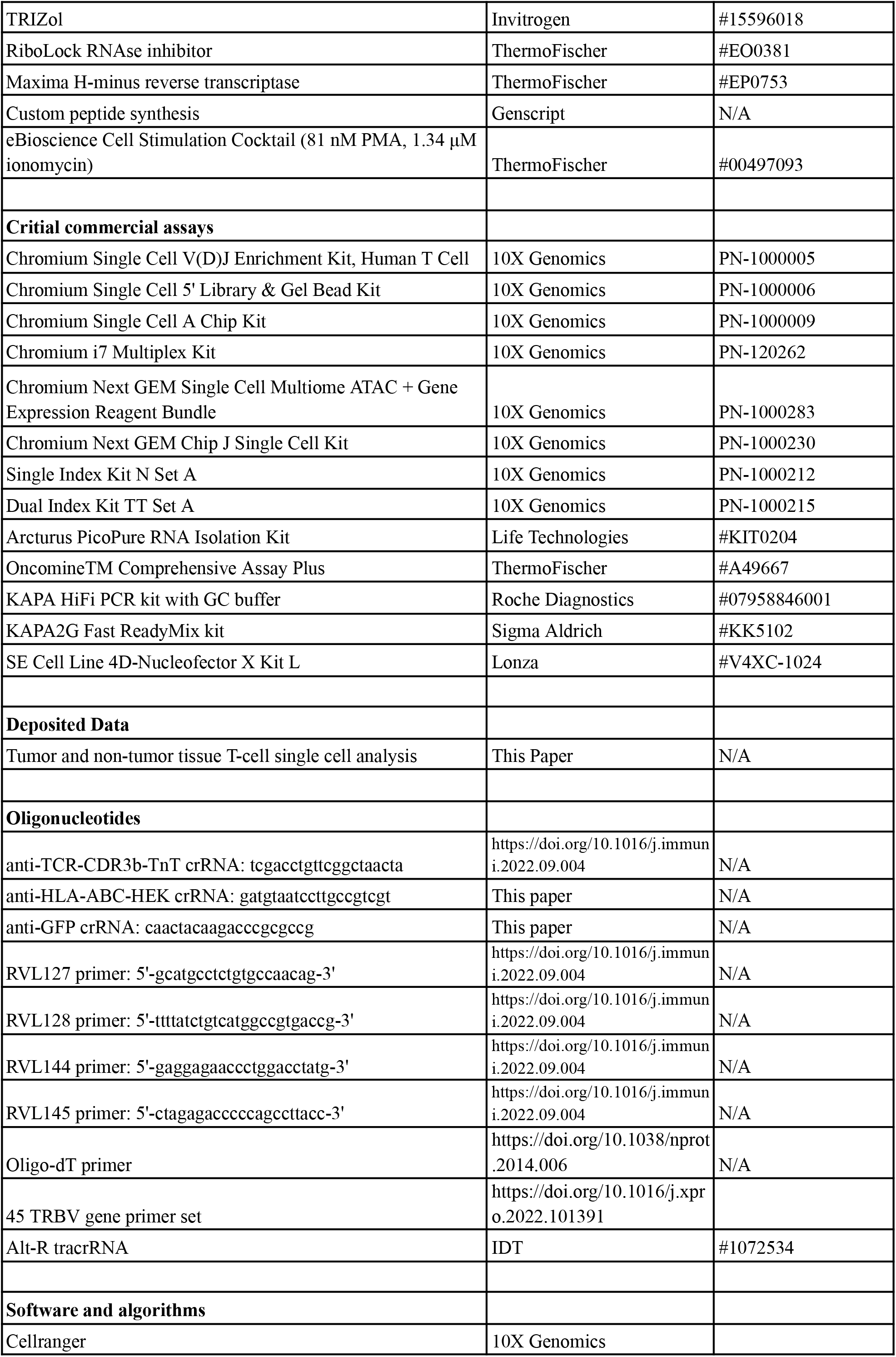
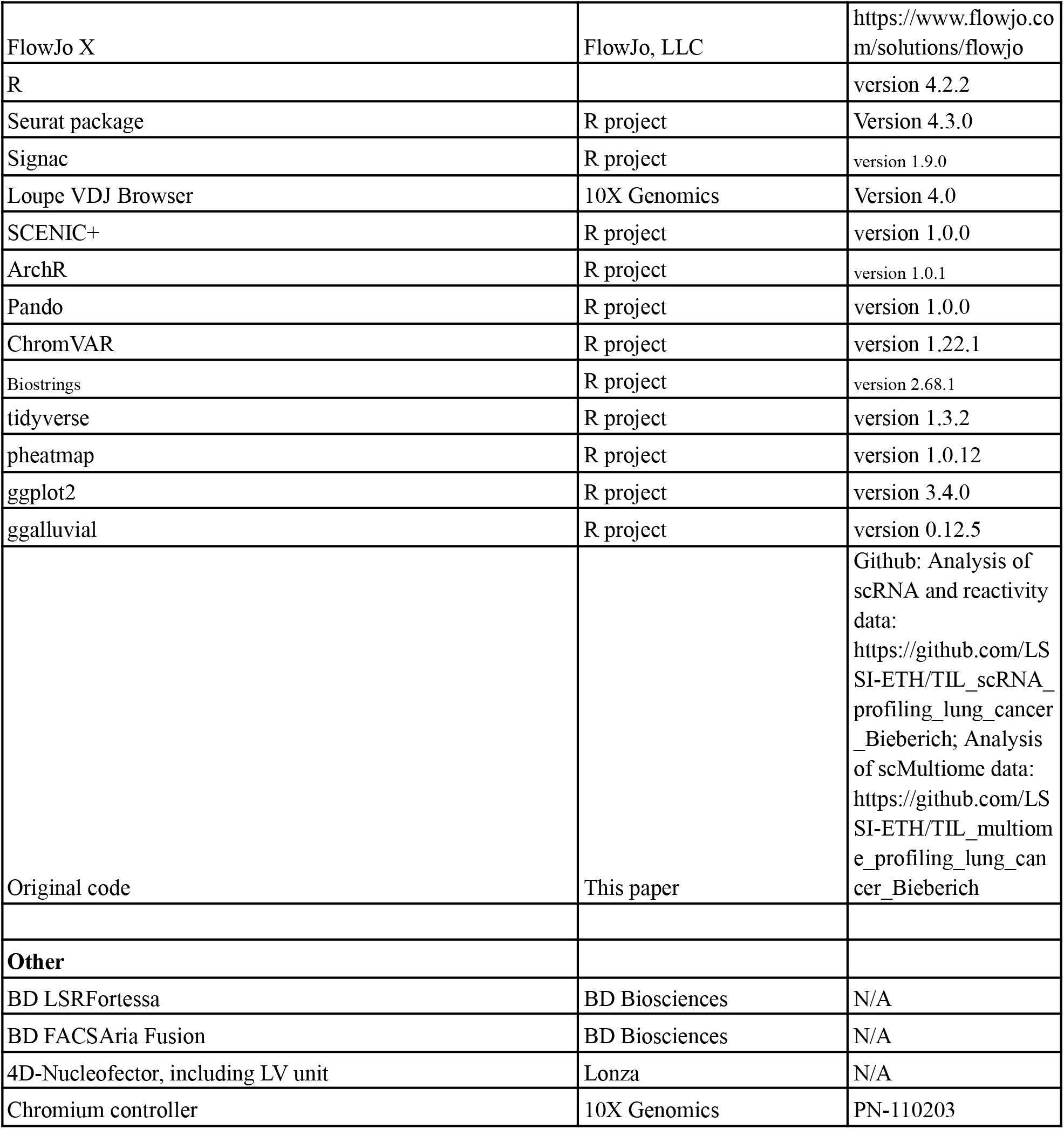

### Patient information and biospecimen processing

Patient samples were collected from individuals with lung cancer that had undergone primary resections and no systemic therapy (i.e. no chemotherapy, targeted therapy or immunotherapy) at Basel University Hospital between 2016 and 2019. The patients described in this study provided written informed consent. All bio-specimens were obtained from patients with stage II-III lung cancer. Detailed patient characteristics are provided in **Table S1**.

Primary resections of lung cancer lesions were immediately processed into single-cell suspensions through mechanical dissociation and enzymatic digestion using accutase, collagenase IV, hyaluronidase and DNAse type IV (Stemcell). Single-cell suspensions of samples were cryopreserved until experimental use.

### HLA and tumor exome sequencing

HLA sequencing of peripheral blood mononuclear cells was performed by the pathology department at Basel University Hospital. Tumor exome sequencing was done using the commercially available OncoPanel platform (OncomineTM Comprehensive Assay Plus - DNA). Sequencing and analysis were performed by the Pathology Unit, led by Matthias S. Matter at Basel University Hospital.

### T cell sorting

Tumor single-cell suspension was thawed, washed, resuspended in blocking solution (Fc-block agent 1:50 in FACS buffer) for 15 min at 4 °C. Staining solution was added and suspension was incubated for 30 min at 4 °C (fluorescently-labeled antibodies were diluted in FACS buffer to 1 ug/mL). Cells were washed twice and cell sorting was performed using a BD FACS Fusion in FACS buffer (DPBS with 1% FBS). The fluorescently-labeled antibody staining consisted of anti-CD8 (BV421, Biolegend), anti-CD4 (APC, Biolegend), anti-CD3 (PE/Cy7, Biolegend), anti-CD45 (PE, Biolegend) and DAPI. TILs (CD8^+^, CD4^-^ and CD3^+^) were immediately subjected to single-cell droplet encapsulation.

### Single-cell sequencing of TILs

Sorted TILs were counted using a hemocytometer and diluted at the desired cell concentration used in for the 10x single-cell sequencing protocol (1,000 cells/uL). The single-cell 5’ VDJ and 5’ GEX kits (10X Genomics) were used to capture transcriptomic information as well as allowing for a target enrichment step to capture immune repertoire information of single-cell paired TCR alpha and beta chains. Encapsulation of lymphocytes and DNA-barcoded gel beads was performed using the Chromium controller (10x Genomics, PN-110203). Briefly, 10’000 - 20’000 cells were loaded per channel for a targeted recovery of 5’000-10’000 droplet encapsulated cells. Reverse transcription and preparation of single-cell transcriptome and TCR libraries was done according to the manufacturer’s instructions (CG000086 Rev M, 10X Genomics) and using the following kits: PN-1000006 and PN-1000005. Libraries were sequenced at 5000 (VDJ), 20000 (Transcriptome) and 25000 (ATAC) paired-end reads per cell.

For single-cell Multiome sequencing library preparation, the following changes were made to the protocol: Sorted TILs were subjected to nuclei isolation according to the manufacturer’s instructions (CG000365). Isolated nuclei were adjusted to the desired concentration (2,500 cells/uL) and incubated in a Transposition Mix before also loading them onto a channel for encapsulation. Preparation of transcriptome and chromatin accessibility libraries were done according to the manufacturer’s protocol (CG000338).

### Single-cell data pre-processing and quality control

CellRanger software version 5 was used to demultiplex fastq reads from transcriptome, ATAC and TCR scSeq libraries and align them to the GRCh38 reference human genome. Subsequent data QC and analysis was performed using R (version 4.2.2) and the Seurat (version 4.3.0) ^94^, Signac (version 1.9.0) ^95^ and ArchR (version 1.0.1) ^96^ packages. QC steps consisted of the exclusion of TCR and BCR genes (prevention of clonotype influence on subsequent clustering), the exclusion of cells with lower than 150 or greater than 4500 genes (low quality cells), and the exclusion of cells in which more than 10% of UMIs were associated with mitochondrial genes (reduction of freeze-thaw metabolic effects).

### Single-cell data integration and clustering

Single-cell RNA sequencing data was read into R using *Read10X*. A Seurat object was generated for each patient (*CreateSeuratObject*), all patients were merged into the same object (*merge*) and QC steps as mentioned before were applied. The Seurat object was split into individual object (by patient) and normalization (*SCTransform*) was applied and mitochondrial genes were regressed out. Variable features were selected from the list of normalized patient Seurat object (*SelectIntegrationFeatures*) and Seurat objects were merged again (*merge*). Previously extracted variable features were set and principal component analysis (PCA) was performed (*RunPCA*) on the normalized assay data. To reduce batch effects, we used the Harmony (version …) package (*RunHarmony*). Finally, UMAP was calculated, and clusters were generated (*RunUMAP*, *FindNeighbors*, *FindClusters*).

### scTCR-seq data analysis

T cell receptor data was analyzed using R Studio and Loupe VDJ Browser 4.0. In brief, TCR information was read into R after Cellranger analysis of the respective fastq files to generate the filtered_contig_annotations.csv file. Using R, clonal definitions were made based on abundance of an exact match of the amino acid CDR3α and -β sequence. Single chains were filtered out and clones that matched with multiple chains (more than one Vα or Vβ) were separated from clones that matched with exactly one Vα and Vβ chain.

### Single-cell multiome analysis

Signac: Multome analysis performed with the Signac package consisted of the following 7 steps: 1. Read in multiome data and create multimodal Seurat object (*Read10x_h5, CreateSeuratObject, CreateChromatinAssay, NucleosomeSignal, TSSEnrichment, CallPeaks, keepStandardChromosomes, subsetByOverlaps, FeatureMatrix, CreateChromatinAssay*); ATAC QC settings as in Signac documentation. 2. Create a common set of peaks (*reduce, width, combined.peaks, FeatureMatrix, CreateChromatinAssay, RunTFIDF, FindTopFeatures, RunSVD*). 3. Merge ATAC (*merge*). 4. Integrate ATAC via harmony (*RunHarmony*). 5. Normalize RNA (*SCTransform*). 6. Integrate SCT via PCA, harmony and UMAP (*SelectIntegrationFeatures, merge, RunPCA, RunHarmony, RunUMAP, FindNeighbors, FindClusters*). 7. Merge ATAC and RNA (*FindMultiModalNeighbors, RunUMAP, FindClusters*).

Single-cell multiome (scRNA and scATAC) inferences for gene regulatory network building are supported by several bioinformatic tools such as SCENIC+, ArchR, Pando and Signac.

ChromVAR, ArchR and SCENIC+ were used according to the official documentation.

### Query mapping of scRNA-seq data onto scMultiome data

Query mapping was performed using the integrated functions of the Signac package. Transfer anchors between reference (multiome data) and query (RNA data) were identified (*FindTransferAnchors*). These were then used to map the query onto the reference locations (*MapQuery*).

### TCR selection and synthesis for reactivity testing

Analysis of the single-cell RNA + TCR sequencing data, allowed for a guided selection of clonotypes. Initially, the top 10 largest clones were already selected for testing. Additional clones were selected using the following criteria: 1. Select barcodes from clusters of interest (here *Exhausted*, *RM_exh1*, *RM_exh2* and *Proliferating*). 2. Check clonal frequency in that cluster subset. 3. Filter clones by applying the following selection criteria: (A) Percentage of clonotype in that cluster needs to be >5%. (B) Select top 10 hits from each cluster subset. Based on these criteria the clones with highest abundance in one of the respective clusters were selected. Further clones were selected that were abundant in both the EM1 as well as in one of the previously mentioned clusters.

Paired TCR sequences were selected from the sequencing information and similarly designed and ordered as mentioned in the TCR-Engine pipeline ^60^. In brief, the sequence design had the following order of components: VDJalpha-TRAC-P2A-VDJbeta-TRBC exon 1 (with splicing domain to splice with the endogenous TRBC exon 2). Sequences were ordered through GenScript.

### CRISPR-based generation of TCR libraries in TnT cells for reactivity testing

We used the same CRISPR-Cas9-based knock-in procedure as mentioned in the TCR-Engine publication. In brief, we PCR amplified the HDR template from the received plasmids using Primers RVL127 and RVL128. We mixed up to 43 different HDR templates together at equimolar concentration of 300 ng per plasmid. We then used the Lonza Nucleofector with the SE kit and the “Jurkat E6.1 new” setting with 1x10^5^ TnT cells, 250 ng of HDR template mix and 0.7 uL of anti-TCR-CDR3b-TnT crRNA. After electroporation, TnT cells were cultured in ATCC-modified RPMI-1640 (Thermo Fisher, #A1049101) medium and HDR enhancer (Alt-R HDR Enhancer, IDT) was added at 1:100 for 16 hours. One week post electroporation, TnT cells were FACS sorted using DAPI and anti-CD3 (PE/Cy7, Biolegend) according to the previously mentioned staining protocol. CD3^+^ TnT cells were cryobanked until further use.

### Peptides for co-culture

Peptides and peptide pools used in targeted reactivity testing, were diluted in DMSO and added to the co-culture setup at a final concentration of 1 ug/mL (EBV and TAA peptide pools, and custom synthesized epitopes) or 2 ug/mL (CMV peptide pool) per peptide according to the manufacturer’s instructions.

### Co-culture and FACS of reactive TnT cells

Target to effector cells were co-cultured for 16 hours at a 1:1 ratio. Anti-CD28 (final concentration 2 ug/mL, Life technologies) and recombinant human IL-2 (final concentration 50 U/mL, Peprotech, #200-02) were added at the start of the co-culture. As positive control, PMA+Ionomycin (1:500, Life technologies) or PHA (10 ug/mL, Invivogen, #inh-phap) were used. Target cells, expressing patient autologous HLA class I’s were generated as described below. Patient material co-culture with TnT cells: One day before co-culture start, tumor suspension and PBMCs were thawed and plated with recombinant human IFNy at 200 ng/mL (Peprotech, #300-02) (only for tumor suspension) overnight in X-vivo 15 medium (Lonza, #02-060Q) and washed in PBS prior to the co-culture start. Co-culture was started by combining TnT cells and tumor suspension/PBMC cells at a ratio of 1:1. For PBMC co-cultures, peptides were added at above mentioned concentrations.

After 16 hours of co-culture, reactive TnT cells were sorted via anti-CD3 (PE/Cy7, Biolegend), anti-CD69 (APC, Biolegend) and NFAT-GFP (staining protocol was performed as mentioned above). Cells were sorted into 200 uL of FACS buffer, supplemented with 800 uL of TRIZOL (Sigma-Aldrich, #T9424) and frozen at -80 °C.

### Library preparation of sorted TnT cells, deep sequencing and data analysis

RNA from sorted and frozen TnT cells in TRIZOL, was extracted using the Arcturus PicoPure RNA Isolation Kit (Life Technologies, #KIT0204), according to the manufacturer’s protocol. RNA quantity was determined using a NanoDrop device (ThermoFischer).

cDNA was generated via reverse transcription of an established protocol used in the TCR-Engine publication. In brief, RNA (200 ng) was mixed with Oligo dT Primer (Picelli et al., SMARTSeq2), dNTPs in a total of 36 uL of water. Mix was incubated for 3 min at 72 °C. Following incubation, Maxima RT (2.5 uL, ThermoFischer, #EP0752), Ribolock (1.5 uL, Thermo Scientific, #EO0381) and 5x RT buffer (10 uL, ThermoFischer, #EP0752) were added and the total mix was incubated for 30 min at 50 °C, then 5 min at 85 °C.

RT-PCR was performed using established primers (RVL144 and RVL145) as described in TCR-Engine, using the KAPA Hifi HotStart ReadyMix (Roche, #KK2602). Annealing temperature was set to 62 °C for 25 cycles.

RT-PCR product was QC’ed using the DNA high sensitivity Bioanalyzer Chip (Agilent Technologies). Sample preparation for Illumina deep sequencing was done using the KAPA HyperPrep Kit, PCR-free (Roche, #KK8503) with KAPA Unique-Dual Indexed (UDI) Adapters (Roche, #8861919702). Sequencing was performed on a MiSeq 250PE flow cell (Illumina).

Libraries were sequenced on a MiSeq sequencer (Illumina) for 2 × 250 cycles using MiSeq PE cluster generation kits and MiSeq SBS Kit sequencing reagents (Illumina).

Data analysis was performed in R using the Biostrings package (version 2.68.1).

### TCR reactivity classification

TCR clone frequencies were compared for all control co-culture conditions (pre co-culture, PBMC co-culture, PHA-activated co-culture) and the highest frequency per TCR selected. To generate a fold-change for each target condition per TCR clone, each TCR clone target condition frequency was compared to the previously determined, highest scoring, control. TCR reactivity threshold was selected based on the value density plot across all TCR-condition fold change values, which showed a separation from the low-fold changes at 2.08 (as can be seen in Fig 2C right). To determine the uncertainty in our deep sequencing results and experimental conditions, we calculated the difference between replicates (i.e. Patient BS833, clone 6, CMV_A and CMV_B) in the log2() space: log2(Fold-change clone 6 CMV_A) - log2(Fold-change clone 6 CMV_B), by doing this for all high scoring replicates, we calculated a mean log2()-based uncertainty value of 0.47 that could be combined with out previously determined threshold of 2.08 using this formula: 2^(log2(2.08) +/- 0.47). This resulted in upper and lower uncertainty fold-change values of 2.9 and 1.5, rendering all values above 2.9 as highly significantly reactive.

TCR clones with a fold-change value of >2.5 were termed reactive by taking the 50% of the uncertainty (2.9-2.08 = 0.82) on top of the chosen threshold of 2.08 into account (2.08+0.82/2 = 2.49).

### Neoantigen identification and synthesis

OncoPanel platform (OncomineTM Comprehensive Assay Plus) - based analysis identified tumor-specific mutations. The surrounding 9 amino acids (AA) on each side (19 AA total) were selected to generate 9 AA long peptides (11 different peptides per mutation). netMHCpan 4.1 was used to screen peptide sequences for binding affinity to autologous HLAs. The upper 30% of peptides, sorted based on HLA-binding affinity, were selected for synthesis by Genscript.

### Generation of patient-specific HLA-expressing cell line

HEK-Blue cells were electroporated with a guide RNA (gRNA) targeting the CCR5 genomic locus in complex with Cas9 recombinant protein (rCas9) and a double-stranded DNA homology-directed repair (HDR) template containing the following elements: (i) left and right homology arms of 924 and 906 bp, respectively mapping to the CCR5 genomic locus; (ii) constitutive chicken beta-actin promoter; (iii) Cas9 encoding gene; (iv) nuclear localization signal; (v) T2A peptide; (vi) puromycin resistance gene; (vii) bGH promoter; (viii) GFP encoding gene. FACS was performed to isolate cells based on GFP expression and a clonal cell line was genotyped and phenotyped to confirm integration and expression of Cas9 from the CCR5 locus. The resulting cell line expressing Cas9 and GFP was subjected to a second step of CRISPR-Cas9 genome editing. Second step, we use a guide RNA (gRNA) targeting the native HLA-ABC genomic locus to knock out the native HLA expression. In a third step of genome editing, we replaced the GFP transgene for a gene encoding the HLA allele of interest. For this purpose, HEK Gen-1 cells were electroporated with a gRNA targeting the GFP transgene located in the CCR5 genomic locus and a double-stranded DNA homology-directed repair (HDR) template containing the following elements: (i) left and right homology arms of 796 and 935 bp (ii) HLA-A*0201, (iii) sv40 terminator. A clonal cell line was genotyped and phenotyped to confirm replacement of GFP with the HLA allele of interest at the CCR5 locus. The resulting cell line expressed Cas9 and the HLA allele of interest.

### TCR amplification, sequencing and analysis from single-cell multiome cDNA

An equimolar primer pool of 45 TRBV gene primers was ordered and generated ^97^. Primer pool and partial read 1 primer (10x Genomics barcode binding) together with KAPA Hifi ReadyMix were used at 59 °C annealing temperature for 35 cycles, according to the described protocol from Hudson et al. Following PCR clean up, library preparation was done as mentioned before. Sequencing analysis was done as previously described for TnT sequencing.

## Data and Code availability

- Raw FASTQ files from deep sequencing that support the findings of this study will be uploaded to SRA (NCBI) and will be publicly available as of the date of publication, see resources table.

- Original code for the single cell RNA, multiome and deep sequencing analysis has been deposited on Github and will be uploaded to Zenodo, see resources table.

- Any additional information required to reanalyze the data reported in this paper is available from the lead contact upon request.

## Acknowledgements

We thank Dr. Jack Kuipers for his advice on the statistical analysis of TCR reactivity deep sequencing data. We thank the ETH Zurich D-BSSE Single-Cell Facility for their assistance with FACS, especially Mariangela Di Tacchio and Chiara Cavallini and the ETH Zurich D-BSSE Genomics Facility Basel for their assistance with single cell and deep sequencing experiments, especially Ina Nissen and Dr. Christian Beisel. This study is supported by funding from the Personalized Health and Related Technologies Grant (to S.T.R.) and the NCCR Molecular Systems Engineering (to S.T.R.).

## Author contributions

F.B., R.V.-L. and S.T.R. designed the study; F.B., R.V.-L., M.T., M.T., M.M., A.Z. and S.T.R. contributed to experimental design; F.B., R.V.-L., K.-L.H., P.H. performed experiments; F.B. and H.J. analyzed data; F.B. and S.T.R. wrote the manuscript with input from all authors.

